# Characterization of the structure and self-assembly of two distinct class IB hydrophobins

**DOI:** 10.1101/2022.08.29.505574

**Authors:** Calem Kenward, Kathleen L. Vergunst, David N. Langelaan

## Abstract

Hydrophobins are small proteins secreted by fungi that accumulate at interfaces, modify surface hydrophobicity, and self-assemble into large amyloid-like structures. These unusual properties make hydrophobins an attractive target for commercial applications as emulsifiers and surface modifying agents. Hydrophobins have diverse sequences and tertiary structures, complicating attempts to characterize how they function. Here we describe the atomic resolution structure of the unusual hydrophobin SLH4 and compare its function to another hydrophobin, SC16. Despite containing only one charged residue, SLH4 has a similar structure to SC16 yet has strikingly different rodlet morphology and propensity to self-assemble. Secondary structure analysis of both SC16 and SLH4 before and after assembly suggest that residues in the first intercysteine loop undergo conformational changes. This work outlines a representative structure for class IB hydrophobins and illustrates how hydrophobin surface properties govern self-assembly, which provides context to rationally select hydrophobins for applications as surface modifiers.

**Keypoints:** -The atomic-resolution structure of the hydrophobin SLH4 was determined using nuclear magnetic resonance spectroscopy

-The structure of SLH4 outlines a representative structure for class IB hydrophobins

-The assembly characteristics of SLH4 and SC16 are strikingly different, outlining how surface properties of hydrophobins influence their function.

## Introduction

Hydrophobins are a diverse group of small (8-14 kDa) proteins produced by Ascomycota and Basidiomycota fungi that readily form ordered assemblies. Fungi secrete hydrophobins into the environment to play key roles in the fungal life cycle: reducing the surface tension of water and allowing hyphae growth into the air, assisting in the attachment of pathogenic fungi to plants, and coating spores to resist wetting and allow proper dispersion (Talbot et al. 1993; Wessels 1994; Talbot et al. 1996; Pham et al. 2016). These unusual proteins are excellent emulsifiers, foam stabilizers, and surface modifiers with their remarkable ability to modify the properties of hydrophobic-hydrophilic interfaces or surfaces (Hektor and Scholtmeijer 2005; Opwis and Gutmann 2011; Green et al. 2013; Ren et al. 2013; Wösten and Scholtmeijer 2015; Pham et al. 2016). Due to these unique properties, hydrophobins are a target of interest for commercialization in pharmaceuticals to improve the bioavailability of water-insoluble drugs (Patravale et al. 2004; Haas Jimoh Akanbi et al. 2010),^9,13^ the food industry as foam stabilizers to increase the longevity of food textures like in ice cream (Wösten and Scholtmeijer 2015), as well as for the production of biosensors and stimulation of tissue growth (Corvis et al. 2005; Zhao et al. 2007; Hou et al. 2008; Li et al. 2009). However, the precise mechanism of hydrophobin self-assembly remains unresolved, thus limiting the ability to rationally design hydrophobins for these applications.

Hydrophobins can be subdivided into two classes based on sequence composition, the intercysteine sequence lengths between their eight conserved cysteine residues, their origin, and their assembly properties (Wessels 1994; Wösten et al. 1994; Wösten and De Vocht 2000; Sunde et al. 2008). Class I hydrophobins are produced by Ascomycetes and Basidiomycetes and form highly durable and insoluble amyloid-like structures called rodlets, which are resistant to denaturation by heat or organic solvents and can efficiently coat surfaces (Hess et al. 1968; Wessels et al. 1991). Class I hydrophobins are highly variable in size, function, and biophysical properties. Sequence alignments and principle component analysis indicate that class I hydrophobins can be further subdivided into those originating from Ascomycota or Basidiomycota fungi (class IA and class IB, respectively), with class IB hydrophobins having more consistent intercysteine sequence lengths (Wösten 2001; Linder et al. 2005; Gandier et al. 2017). In contrast to class I hydrophobins, class II hydrophobins are only produced by Ascomycetes, are smaller (<10 kDa), have greater sequence conservation, and form less robust assemblies at interfaces (Linder 2009; Gandier et al. 2017).

Mirroring the diversity observed in residue composition, the tertiary structures of hydrophobins are diverse (Bayry et al. 2012). A key feature of hydrophobins is the presence of eight cysteine residues that form a conserved pattern of disulfide bonds (Sunde et al. 2008). However, despite this shared trait the intercysteine sequences that connect the β-strands (L_1_–L_3_, also referred to as the C3-C4, C4-C5, and C7-C8 intercysteine loops) often vary in sequence, length, structure, and dynamics. Recently, the first atomic-resolution structure of a class IB hydrophobin was determined for SC16 (derived from *Schizophyllum commune*) (Gandier et al. 2017); however, it is unclear how consistent the class IB hydrophobin fold is, or how hydrophobin sequence properties may correlate with their structure and function.

To guide hydrophobin commercialization and better characterize the correlation between hydrophobin sequence, structure, and function, we compared the solution-state structure, rodlet features, and preferred assembly conditions of two class IB hydrophobins, SC16 and SLH4 (derived from *Serpula lacrymans*). SLH4 is an especially unusual hydrophobin because it only has one charged residue in its sequence (Table S1), which may give it distinct properties from SC16.

## Materials and Methods

### Expression and purification of hydrophobins

Plasmids encoding for hydrophobins with their predicted N-terminal signal peptides removed (SC16: Thr18–Leu114 of the *hyd1* gene of *S. commune*, NCBI ID: EFI94929; and, SLH4: Gly19–Leu104 of the *slh4* gene from *S. lacrymans*, NCBI ID: EGO29586) were the same as those used previously to express and purify class IB hydrophobins (Kenward et al. 2020). Each hydrophobin was produced as a fusion to an N-terminal hexahistidine tag (H_6_) with a tobacco etch virus (TEV) protease recognition sequence (H_6_-SC16 and H_6_-SLH4), and SC16 was also fused to upstream sequences coding for a hexahistidine tag, the B1 domain of protein G, and a thrombin recognition sequence (H_6_-GB1-SC16). The expression, purification, tag removal, and refolding of these hydrophobins was carried out as outlined previously (Kenward et al. 2020). After purification hydrophobins were stored at -20 °C in solution or as lyophilized powder.

### Emulsification Assay

Lyophilized hydrophobins or BSA were resuspended to 100 µg/mL in 2 mL of water containing 25 µg/mL Remazol Brilliant Blue for contrast. Samples were then vortexed with an equal amount of mineral oil for 3 minutes. Images of the samples were acquired pre-agitation and one minute post-agitation.

### Electron microscopy

Lyophilized hydrophobins were resuspended in water at 100 µg/mL. A single drop of hydrophobin solution was deposited onto parafilm and allowed to equilibrate for 20 minutes to allow self-assembly to occur at the air-water interface of the drop. An electron microscopy grid (Cu 200 mesh, formvar/carbon coated) was floated onto the drop for 30 seconds, stained with 2% uranyl acetate for 10 minutes, and then washed with water. Images were acquired using a JEOL JEM 1230 Transmission Electron Microscope with an ORCA-HR Hamamatsu digital camera at the Dalhousie University Electron Microscopy CORE Facility.

### Atomic force microscopy

Hydrophobin samples were prepared in dH_2_O (5 μg/mL). A 50 μL drop of solution was placed onto freshly cleaved highly oriented pyrolytic graphene (HOPG) discs (SPI Supplies® Grade SPI-2) and allowed to dry overnight at room temperature. Intermittent contact mode images were collected using a NanoWizard II Ultra (Bruker) using Tap300AI-G tips. Images were processed using the associated processing software. For each hydrophobin two samples were prepared with 3 images collected per sample.

### Circular dichroism spectroscopy

Hydrophobin samples were prepared at 50 μM in dH_2_O. Unmixed samples were incubated overnight at room temperature. Mixed samples (1 mL) were gently agitated end-over-end overnight with SLH4 vortexed for a further 2 h, followed by 3 h of settling time to allow foam to separate from solution. Spectra were collected as 5x replicates at 20 °C from 180-279 nm using a 0.1 mm cuvette on an Olis DSM20 circular dichroism spectrophotometer. Data were processed in Olis Spectral Works (v. 5.888.272) and MSExcel following the DichroWeb User Guide (http://dichroweb.cryst.bbk.ac.uk/html/userguide.shtml). The BeStSel webserver was used to estimate the secondary structure composition of each sample (Micsonai et al. 2018).

### Thioflavin-T assays

Reactions were set up in a matrix of buffer conditions consisting of a multicomponent buffer (20 mM MES, 20 mM TRIS, 20 mM phosphate) at varying pH (5.5, 6.5, 7.5, 8.5) versus varying [NaCl] (10 mM, 250 mM, 1 M), and contained 50 μg/mL hydrophobin and 20 μM fresh, filtered ThioflavinT (ThT). Reactions were 100 μL and done as 6x replicates in a sealed 96-well black-bottomed plate. A SpecraMax M3 plate reader was used to monitor ThT fluorescence. Every 2 min reactions were shaken for 30 s then fluorescence was measured with λ_ex_ = 430 nm, λ_cutoff_ = 455 nm, and λ_em_ = 480 nm.

### Nuclear magnetic resonance spectroscopy and structure calculation

Lyophilized ^13^C/^15^N-labelled SLH4 (800 µM) was resuspended in NMR buffer containing 20 mM MES pH 6.5, 50 mM NaCl, 1 mM NaN_3_, and 10% D_2_O. Sample quality was assessed by collecting ^1^H-^15^N heteronuclear single quantum coherence (HSQC) spectra and standard triple resonance experiments were analyzed to assign the backbone and sidechain resonances. Distance restraints were derived from ^15^N-edited NOESY-HSQC and both aliphatic and aromatic ^13^C-edited NOESY-HSQC spectra, with mixing times of 110 ms. ^1^H-^15^N heteronuclear NOE experiments were acquired with saturated and reference spectra collected in an interleaved manner with a saturation time of 4 seconds. Spectra were processed using a combination of NMRPipe version 10.6 (Delaglio et al., 1995) and Bruker Topspin version 4.0.6, and analysed using CcpNmr Analysis version 2.4 (Vranken et al., 2005). Experiments were collected on a 600 MHz Varian INOVA NMR spectrometer equipped with a triple resonance probe (Queen’s University, Kingston, ON), on a 500 MHz Bruker Avance spectrometer with BBFO SmartProbe (NMR-3, Dalhousie University), and a 700 MHz Bruker Avance III NMR spectrometer equipped with a TCI cryoprobe (National Research Council, Halifax NS).

Backbone and side-chain ^1^H, ^13^C, and ^15^N chemical shift values, manually assigned NOESY peak lists, and dihedral angle restraints generated by DANGLE (Cheung et al. 2010) were used as inputs to ARIA version 2.3 (Rieping et al. 2007) for automated NOE assignment and structure calculation. Preliminary structures and the chemical shifts of cysteine C^β^ nuclei supported the presence of disulfide bonds, so disulfide bond restraints were incorporated into the structure calculation. Hydrogen bond restraints were added based on preliminary structures for residues that had strong signals remaining in a ^1^H-^15^N HSQC after exchange with D_2_O for 1 hour. Structures were calculated in eight iterations followed by a final water refinement step of the 20 lowest energy structures out of 100 performed by CNS version 1.21 (Brunger 2007). The quality of the final ensemble of structures was assessed through Recall Precision and F-factor analysis (Huang et al. 2005), PROCHECK (Laskowski et al. 1996), and the protein structure validation suite (Bhattacharya et al. 2007). The PyMOL Molecular Graphics System version 2.0 (Schrödinger, LLC) was used for visualization and figure generation. The protein-sol webserver (Hebditch and Warwicker 2019) was used to visualize SLH4 surface hydrophobicity. Chemical shifts and the final 20-member ensemble of structures for SLH4 have been deposited into the Protein Data Bank (PDB) and the Biological Magnetic Resonance Data Bank (Accession numbers 5W0Y and 30301, respectively).

## Results

The SHuffle T7 express strain of *E. coli* was used to overexpress H_6_-SC16, H_6_-SLH4, and H_6_-GB1-SC16. After Ni^2+^ affinity chromatography and subsequent removal of the expression tag using TEV or thrombin protease, reverse-phase high performance liquid chromatography or size exclusion chromatography was used to generate moderate yields (4-12 mg per litre of culture) of highly pure hydrophobins that were suitable for further characterization (Kenward et al. 2020). Both SC16 and SLH4 were able to stabilize oil-water emulsions (Fig. S1), which is a common property of hydrophobins and indicates that both proteins are functional.

### Rodlets formed by SC16 and SLH4 have distinct morphologies

Negative staining transmission electron microscopy (TEM) of grids prepared with non-agitated samples of either SC16 or SLH4 consistently resulted in no visible protein deposition. However, if the TEM grid was floated on top of a drop of solution containing hydrophobins, repeating patterns of assembled hydrophobins were visible, suggesting that both SC16 and SLH4 assemble at the air-water interface (Fig. 1). SC16 appears as large, irregular plaques, often over 100 nm long, that have striations which suggest there is significant internal order within these structures. In contrast, SLH4 assembled into smaller structures (∼20 nm across) that have less pronounced striations. To further characterize these hydrophobin assemblies, we used atomic force microscopy to observe rodlet morphology (Fig. 2). SC16 readily self-assembles and coats HOPG surfaces when dried onto a surface overnight. The SC16 rodlets often formed in pairs, with a paired width of ∼200 Å and lengths that varied from 100 – 500 nm. SC16 rodlets would also efficiently align with each other and form patches of aligned rodlets. In contrast, despite efficiently coating the HOPG surface, SLH4 rodlets were much shorter (20 – 200 nm) and did not align with each other nearly as well as SC16.

**Fig. 1.**
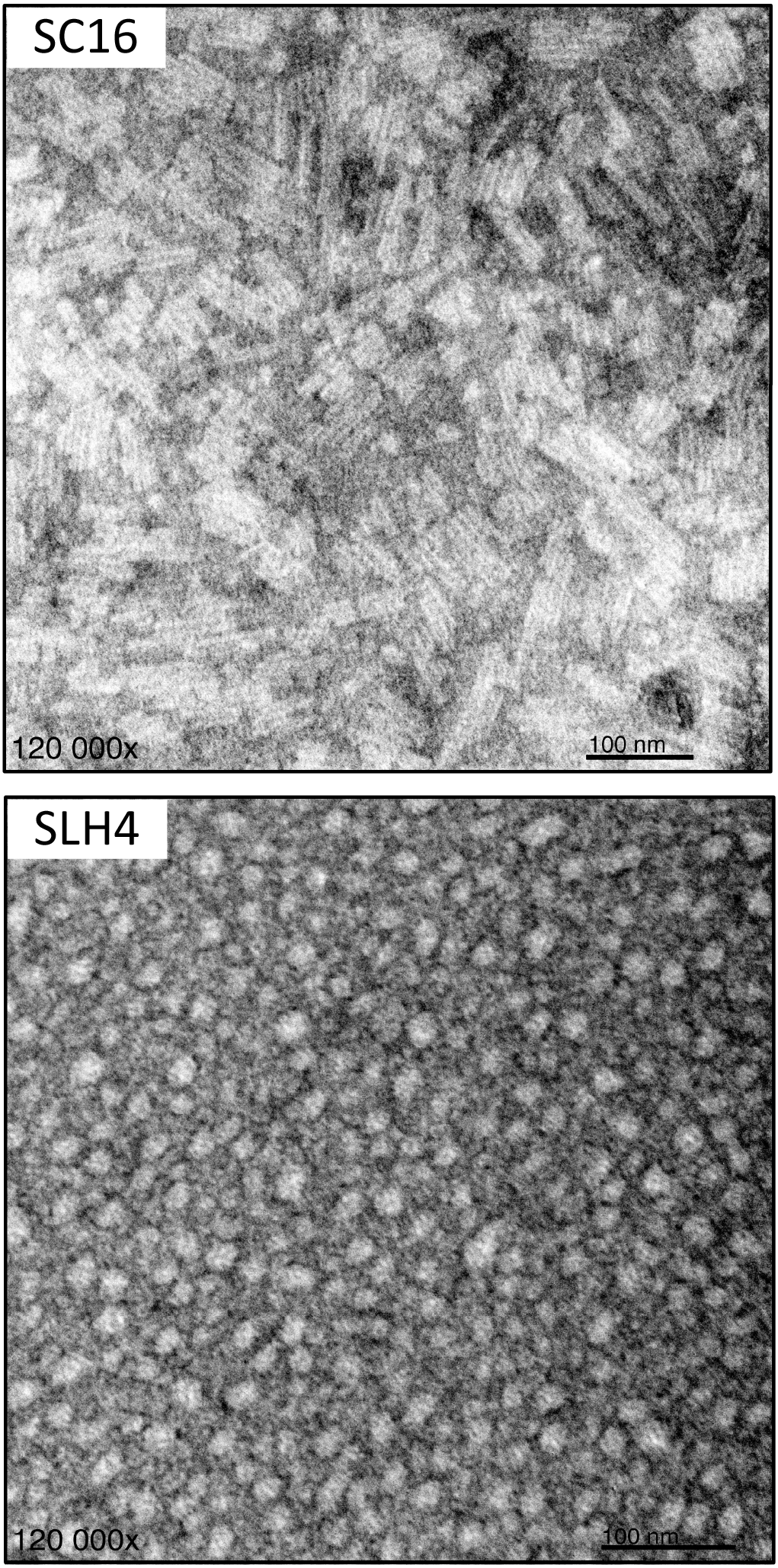
SC16 and SLH4 self-assemble into rodlets at air-water interfaces. Hydrophobins were deposited onto a carbon-coated EM grid, stained with uranyl acetate, and visualized using transmission electron microscopy. Representative fields of view are shown for negatively stained assemblies of SC16 and SLH4.

**Fig. 2.**
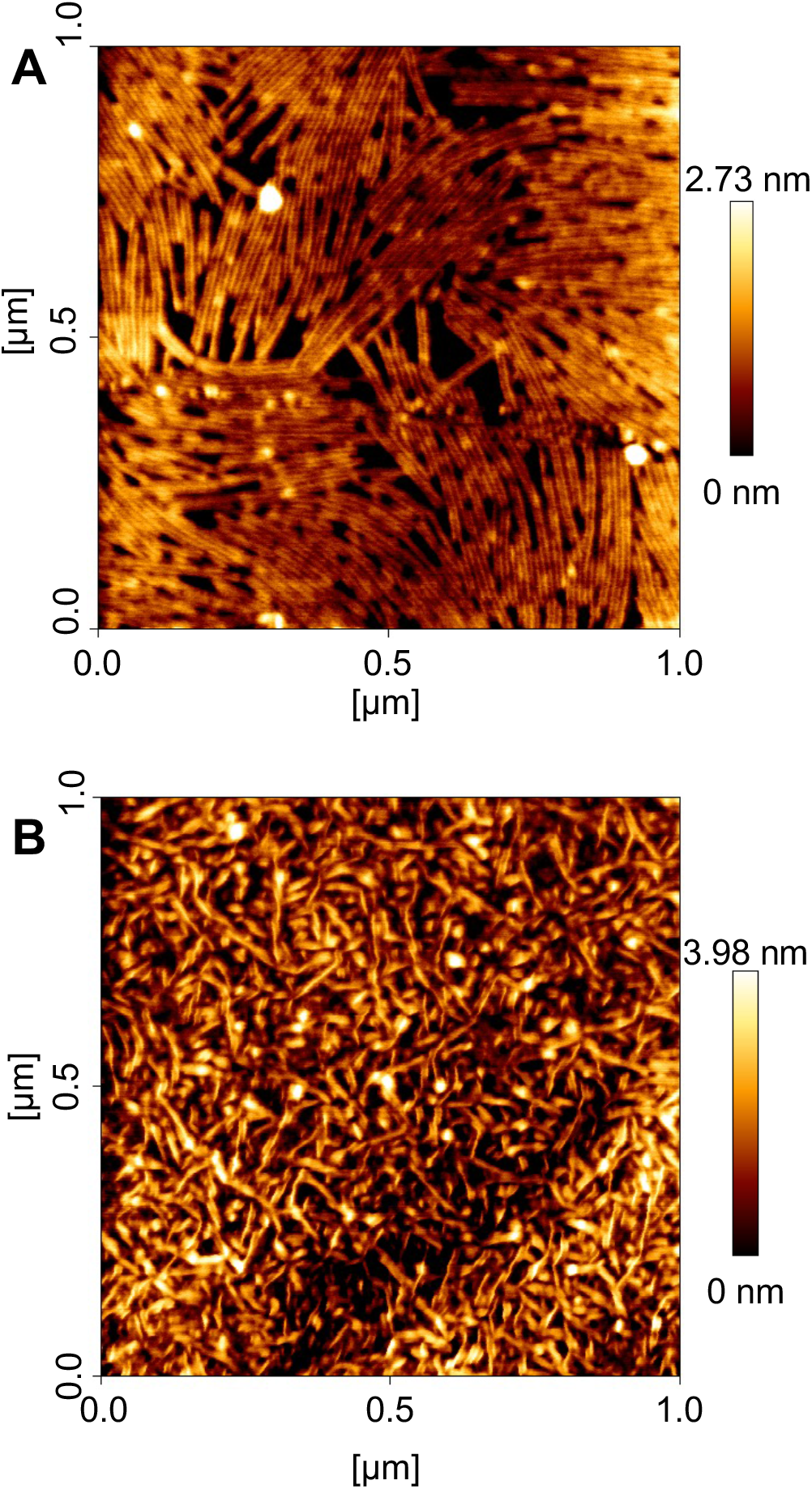
AFM visualization of SC16 and SLH4 rodlets. Hydrophobin samples were incubated on freshly cleaved highly oriented pyrolytic graphite at room temperature and dried overnight. Representative tapping-mode atomic force microscopy images are shown for SC16 **(A)** and SLH4 **(B)**. SC16 rodlets were observed to associate laterally into bundles while SLH4 associated into rodlets that were shorter and less organized.

### SC16 and SLH4 undergo conformational changes during self-assembly

The secondary structure of SC16 and SLH4 both before and after agitation was assessed by circular dichroism spectroscopy (Fig. 3). In the unassembled state both SC16 and SLH4 primarily consist of β-sheet secondary structure with ∼10% α-helix. Upon introduction of an air-water interface through gentle overnight agitation or vigorous vortexing, the solutions of SC16 and SLH4 separated out into a foam that, based on UV-absorbance measurements of the remaining solution, contained all the hydrophobin. Circular dichroism measurements of the foam indicate that there was no α-helix present, and the proportion of total β-sheet increased to ∼37.0% and ∼62% for SC16 and SLH4, respectively.

**Fig. 3.**
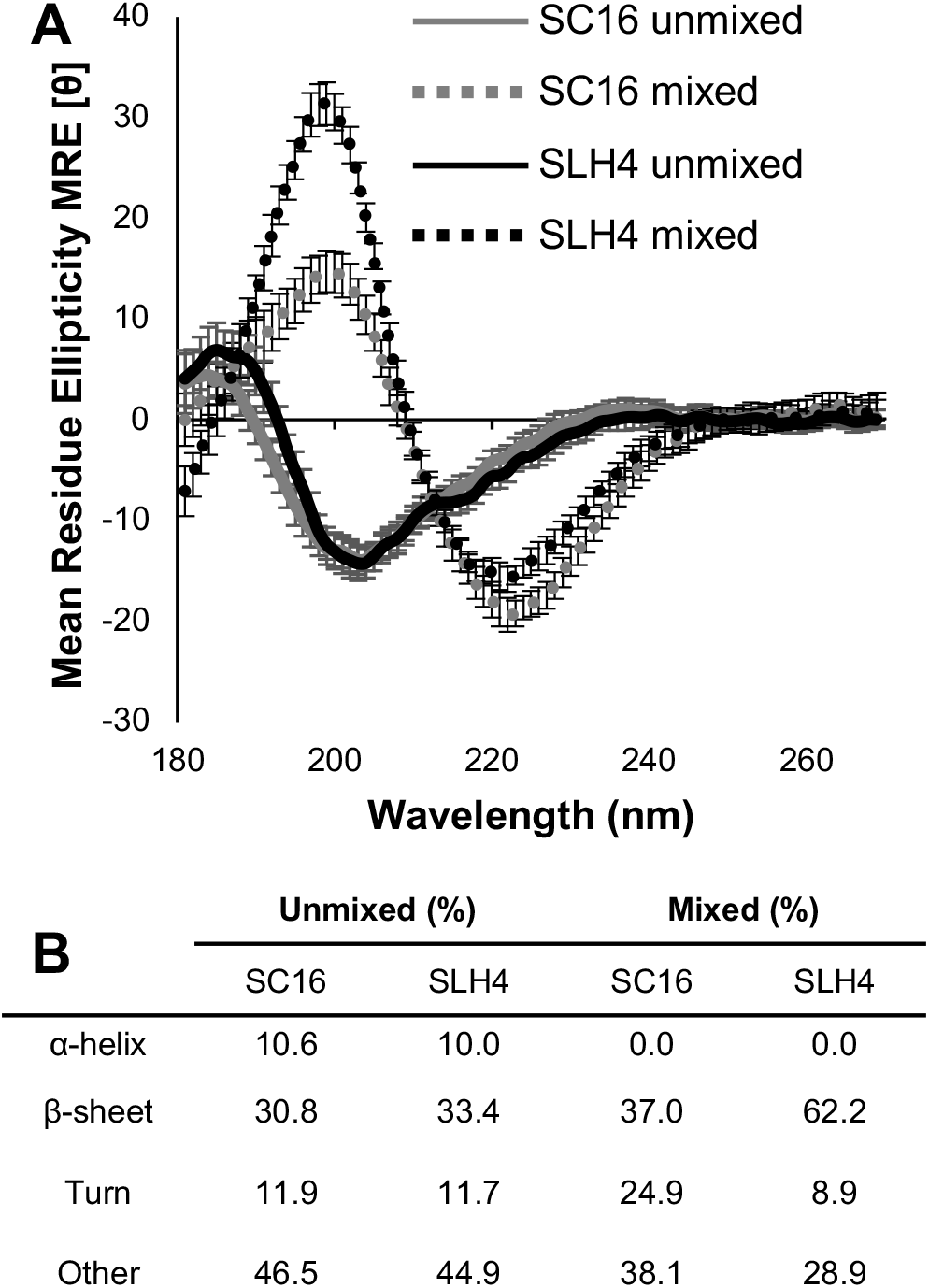
The secondary structure of SC16 and SLH4 changes upon assembly into rodlets. **(A)** Circular dichroism spectra were collected for the unmixed hydrophobin solution and the assembled foam of SC16 (grey) and SLH4 (black) to observe the conformational change undergone during self-assembly due to agitation (solid line: unmixed; dashed line: mixed). Plots show the mean ± S.D. of 5 replicate experiments. **(B)** BeStSel webserver was used to estimate the secondary structure composition of each sample upon agitation (Micsonai et al. 2018).

### Self-assembly of SC16 and SLH4 is influenced by ionic strength and pH

The ability of SC16 and SLH4 to self-assemble in different buffer conditions was quantified using ThT (Fig. 4), which binds amyloids and is frequently used to monitor assembly of hydrophobins and other amyloidogenic proteins (Mackay et al. 2001; Zykwinska et al. 2014; Lo et al. 2014; Gandier et al. 2017). Upon agitation SC16 was readily able to assemble in all conditions tested in the ThT assay, with the fluorescence plateauing within 1 hour (Fig. 4A,B). The degree of assembly increased moderately with an increase in ionic strength or pH of the solution. The ThT assay indicated that SLH4 assembly is much slower than SC16. In contrast to SC16, SLH4 assembly was extremely dependent on buffer conditions, with pH 6.5, 10 mM NaCl being the only combination that allowed for complete assembly within 4 h (Figs. S2 and 4C,D). Based on ThT fluorescence, SLH4 has a much lower ability to self-assemble into amyloid containing rodlets than SC16 and is much more sensitive to the ionic strength of the solution.

**Fig. 4.**
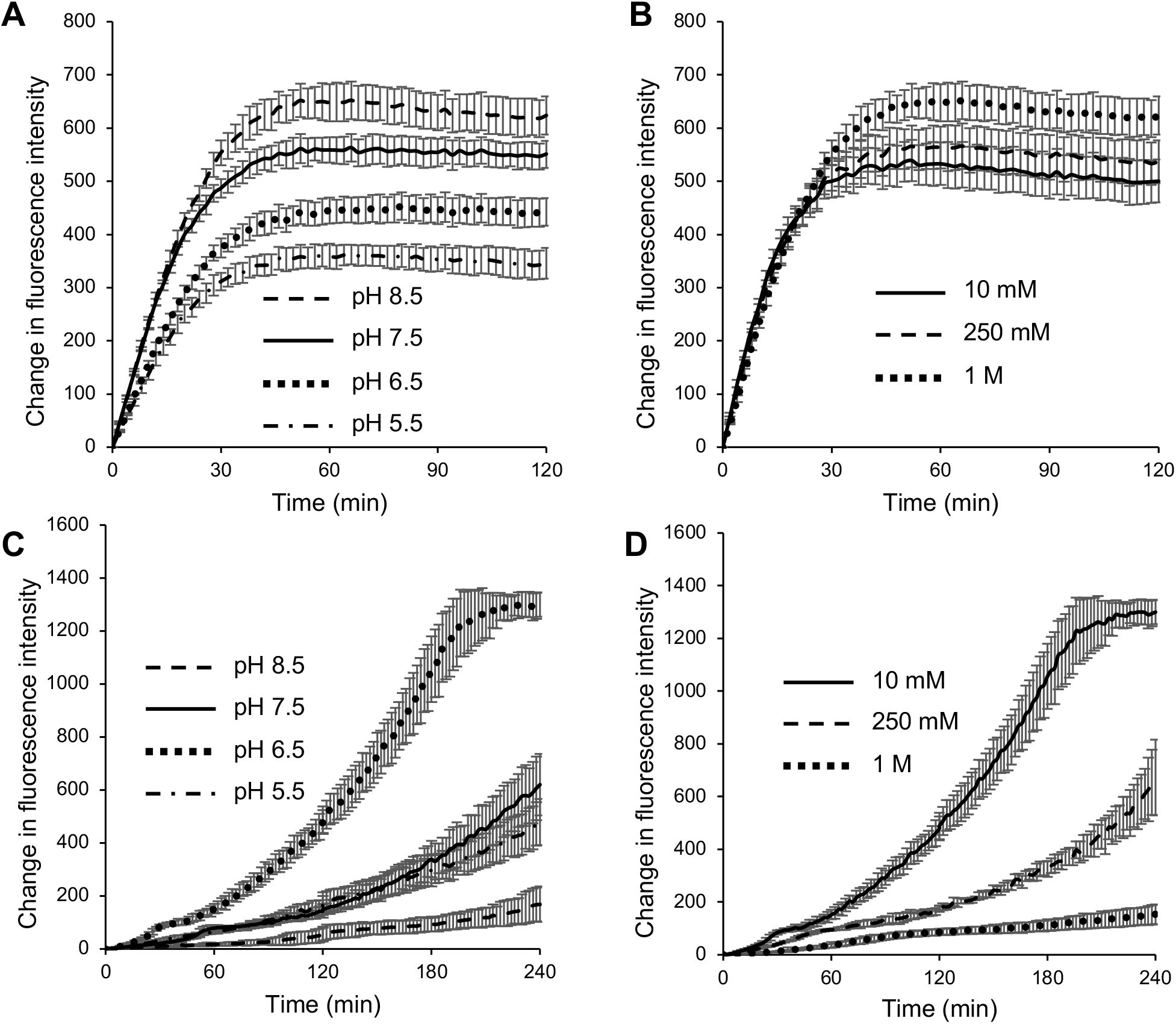
SC16 and SLH4 self-assembly is affected by pH and salt. Thioflavin-T based fluorescence assays were used to quantify rodlet assembly of SC16 **(A**,**B)** and SLH4 **(C**,**D)**. Assembly was monitored in solutions with varying pH (with 1 M or 10 mM NaCl, for SC16 and SLH4, respectively), and in solution with varying [NaCl] (at pH 8.5 and 6.5 for SC16 and SLH4, respectively). Thioflavin-T fluorescence intensity was monitored at 2 min intervals with 30 s shaking between each interval. Plots show mean ± S.D. of 6 replicate experiments.

### The solution-state structure of SLH4

To investigate the mechanistic basis of class IB hydrophobin properties, we used solution state NMR spectroscopy to investigate the structure and dynamics of SLH4 prior to rodlet assembly. The ^1^H-^15^N HSQC spectrum of SLH4 contains signals of relatively uniform intensity with excellent peak dispersion, suggesting that it adopts a stable tertiary structure (Fig. S3). A backbone walk approach was used to assign 99% of backbone and 82% of sidechain resonances. ^15^N-edited and ^13^C-edited HSQC-NOESY spectra were assigned and resulted in 1583 NOE-derived distance restraints for SLH4 (Table 1). The NOE-derived distance restraints as well as dihedral angle and hydrogen bond restraints were used by ARIA to generate a final ensemble of 20 structures for SLH4 (Table 1, Fig. 5) that has excellent conformational statistics, very few distance violations, and converges with a pairwise root mean squared deviation of 0.3 Å for main chain atoms.

**Table 1.**
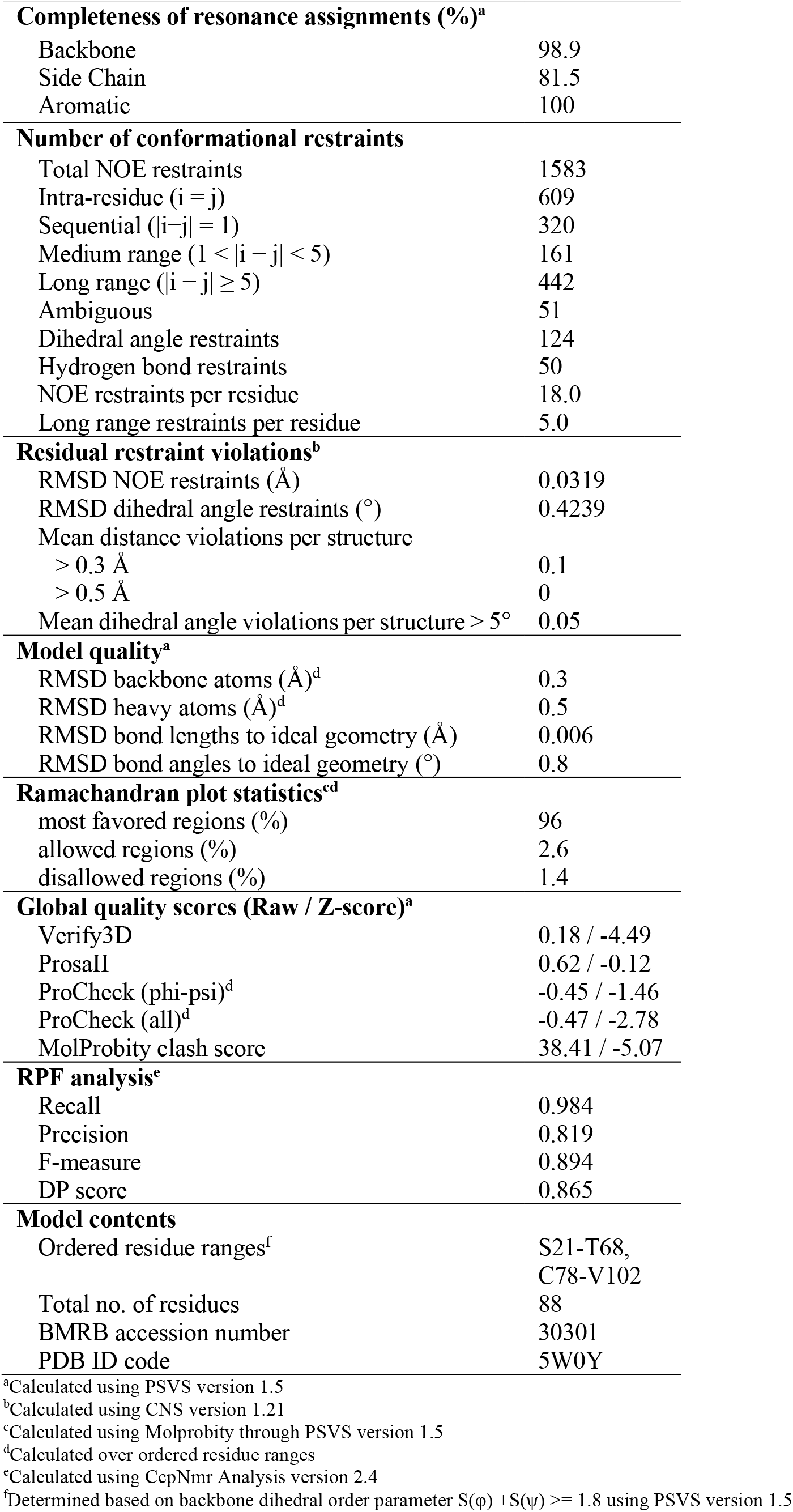
Statistics of the NMR-derived ensemble of SLH4 structures.

**Fig. 5.**
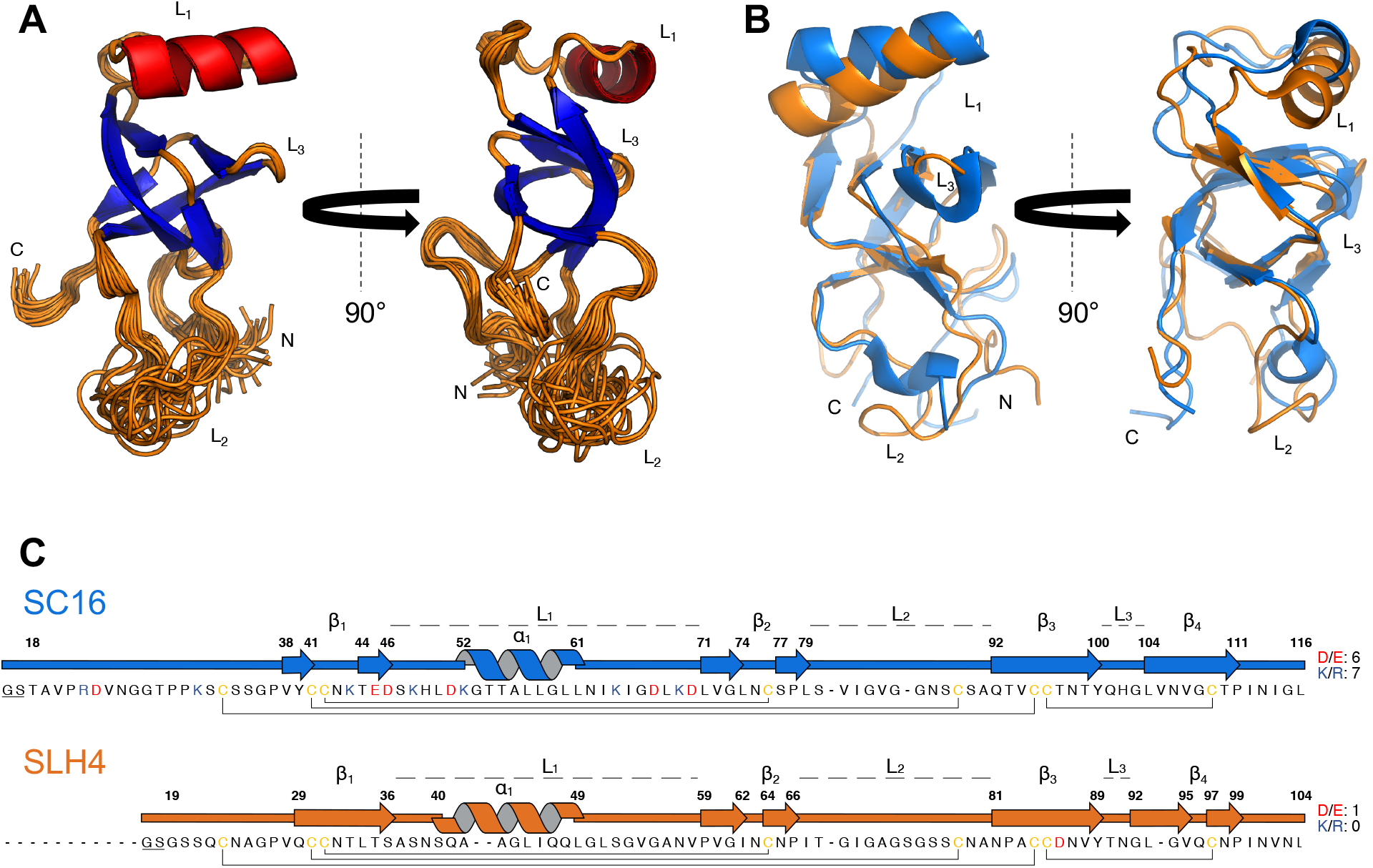
Overview of NMR-derived hydrophobin structural ensembles. **(A)** Ribbon representations of the final 20 lowest energy structures of SLH4 are shown with β-strands and α-helices coloured blue and red, respectively. **(B)** A superposition of the lowest energy structures of SLH4 (yellow) and SC16 (blue; PDB ID: 2NBH; Gandier et al., 2017). Labels denote the intercysteine loops as well as the N- and C-termini of each protein. **(C)** A sequence alignment of SC16 and SLH4 is shown, with acidic, basic, and cysteine residues coloured red, blue, and yellow, respectively. The total number of acidic (D/E) and basic (K/R) residues are listed to the right of the sequence. The secondary structure and regions corresponding to the intercysteine loops (L_1_-L_3_) and β-barrel core (β_1_-β_4_) are indicated. Disulfide bonds are indicated by lines that connect cysteine residues. Residues that are underlined are remnants of protease cleavage and not native to natural hydrophobin sequences.

SLH4 contains β-strands that assemble to form a distorted β-barrel with a hydrophobic core (β_1_: Gln29-Ser36, β_2_: Pro59-Pro66, β_3_: Asn81-Tyr89, β_4_: Gly92-Pro99). β_1_ and β_2_ form an anti-parallel β-sheet with an intervening loop (L_1_; C3-C4) that contains an α-helix from residues Ser40-Leu49. β_3_ and β_4_ form a second antiparallel β-sheet that is separated by a short intervening loop (L_3_; C7-C8). Finally, another loop (L_2_; C4-C5) connects β_2_ to β_3_. The L_2_ of SLH4 has no defined secondary structure, and L_3_ is comprised of a short β-turn and is well-converged in the SLH4 structural ensemble. The N- and C-termini, as well as L_2_, are not as well converged as the rest of the ensemble. This is consistent with heteronuclear NOE measurements (Fig. S4), in that these regions of SLH4 have a reduced value of the heteronuclear NOE (0.1-0.7) compared to the rest of the protein (>0.7), suggesting that these regions are more dynamic and undergo molecular motion on the ps-ns timescale.

Class I and class II hydrophobins often have amphipathic character with pronounced hydrophobic patches on the protein surface (Hakanpää et al. 2006; Kwan et al. 2006; Morris et al. 2013; Ren et al. 2014), which is important for hydrophobin function, as this enhances their accumulation at hydrophobic-hydrophilic interfaces. We used the Protein-sol webserver (Hebditch and Warwicker 2019) to evaluate amphipathic character of SC16 and SLH4 (Fig. 6). SLH4 is hydrophobic, with a polar/non-polar ratio of ∼0.5 over much of its surface. In contrast, SC16 has a single hydrophobic patch localized largely to the side of the β-barrel.

**Fig. 6.**
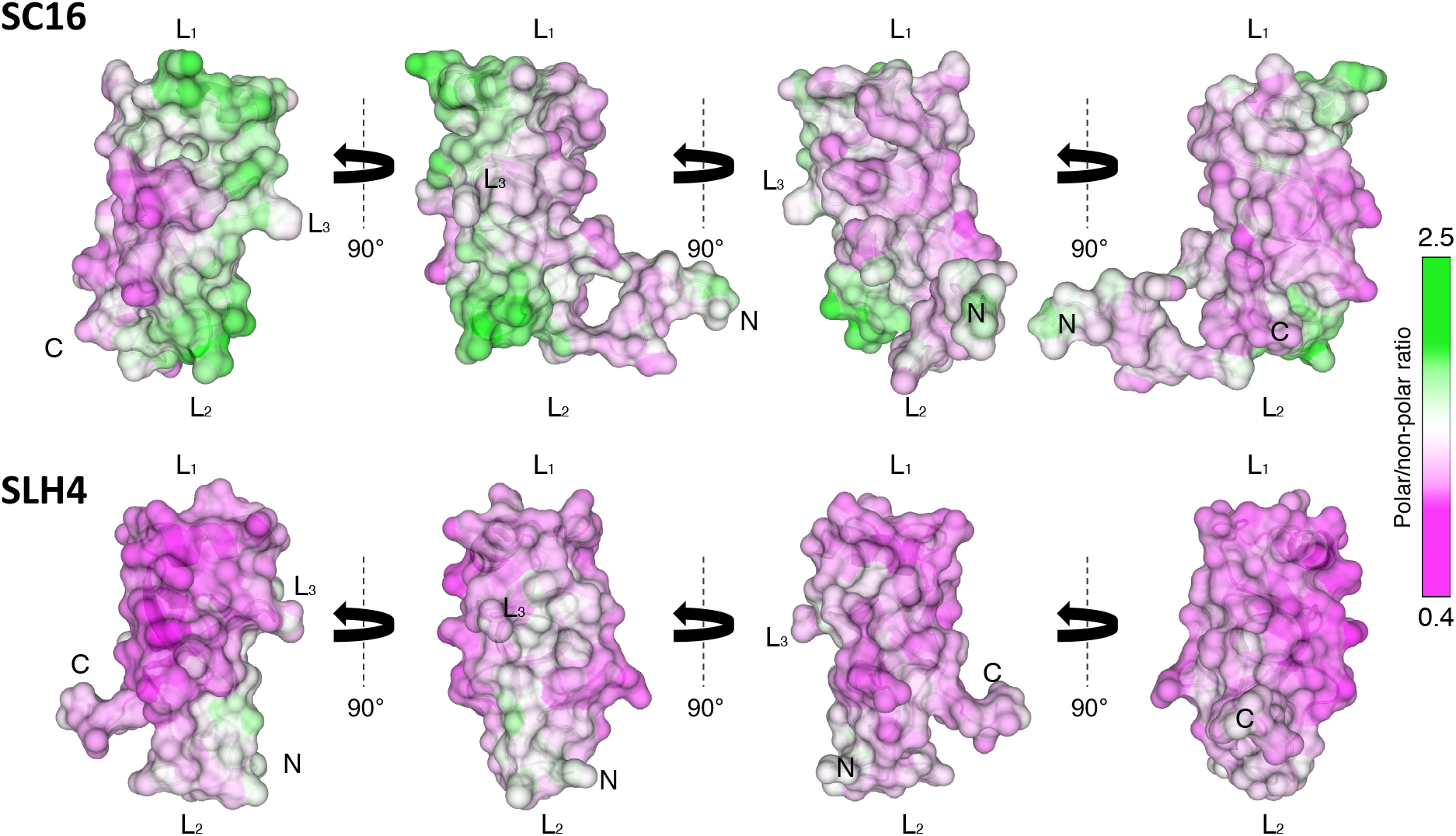
SC16 and SLH4 have distinctive surface hydrophobicity. Surface representations of surface hydrophobicity are shown for SC16 and SLH4. The polar to non-polar ratio of the protein surface was calculated using the protein-sol webserver (Hebditch and Warwicker, 2019), and mapped onto the surface of SC16 (PDB ID: 2NBH) and SLH4.

## Discussion

To aid in the commercial development of hydrophobins, this work investigates the interplay between hydrophobin sequence, structure, and rodlet morphology. Prior to this study, SC16 was the only hydrophobin originating from Basidiomycota that has been structurally characterized and the structural conservation of this family of proteins was unclear (Gandier et al. 2017). The structure of SLH4 that we determined is similar to SC16 (RMSD 2.2 Å; Fig. 5), which suggests a common set of structural features are shared among class IB hydrophobins. This includes a compact globular structure that is centered around several β-sheets that form a distorted β-barrel, and conserved disulfide bonds (consistent with other class I and class II hydrophobins), that covalently link β_1_ to β_2_, β_3_ to β_4_, the N-terminal tail to β_3_, and β_1_ to L_2_. Within L_1_, an α-helix is present that associates with the β-barrel core. Finally, in class IB hydrophobins L_2_ is consistently unstructured and L_3_ forms a β-hairpin that is well-converged in the structural ensembles.

Current models of hydrophobin self-assembly involve significant structural rearrangements of the flexible loops and the formation of an amyloid-like cross-β structure (Macindoe et al. 2012). However, the variability of structural features among class I hydrophobins makes it difficult to define rules for which regions or structures are important for self-assembly. For the class IA hydrophobin EAS (derived from *Neurospora crassa*), deletion and point mutants indicate that L_1_ is not directly involved in rodlet formation, while amyloidogenic sequences within L_3_ are necessary for the formation of rodlets (Kwan et al. 2008; Macindoe et al. 2012). Many other class I hydrophobins contain a sequence that is predicted to be amyloidogenic, but the relative position of this sequence varies: in DewA (derived from *Aspergillus nidulans*) it was predicted to be present in L_2_ (Morris et al. 2013), while in RodA amyloidogenic sequences in both L_2_ and L_3_ were found to be important for self-assembly (Valsecchi et al. 2019).

Together, the features of SC16 and SLH4 provide clues into the mechanism of class IB hydrophobin rodlet formation. In both proteins L_3_ is unlikely to undergo structural changes and participate in amyloidogenesis as it is very short, not dynamic, and is incorporated into the β-barrel core of the protein. Meanwhile, L_2_, while often dynamic, is unlikely to be able to undergo large conformational changes because it is tethered to the β-barrel through the C2-C5 disulfide bond. The consensus amyloid prediction server AmylPred2 (Tsolis et al. 2013) indicates that amyloidogenic sequences are present in L_1_ (Ala56-Asn62 and Leu45-Ile46 for SC16 and SLH4, respectively), which all localize to the α-helix. In both SC16 and SLH4 the interface between L_1_ and the β-barrel is hydrophobic and any movement of L_1_ would present a large hydrophobic surface that could drive accumulation at hydrophobic-hydrophilic interfaces and/or rodlet assembly. A structural change in L_1_ during rodlet assembly is consistent with our circular dichroism spectra that detected loss of α-helical and an increase in β-sheet secondary structure upon agitation. This assembly model is consistent with movements within L_1_ that have been linked to rodlet formation in the related class IB hydrophobin SC3, where protease digestion and hydrogen–deuterium exchange experiments suggest that L_1_ undergoes conformational changes upon binding to interfaces (Wang et al. 2004).

Despite having similar tertiary structures in solution, the assembly propensity and rodlet features of SC16 and SLH4 differ markedly, suggesting that surface properties drive hydrophobin assembly. Surprisingly SLH4 did not assemble as quickly as SC16, despite having a more hydrophobic surface over much of the protein. Despite not assembling as quickly, SLH4 plateaued with higher ThT fluorescence, suggesting that it was able to eventually bind more ThT. Hydrophobic patches are commonly found in hydrophobins and are important for rodlet formation (Hakanpää et al. 2006; Kwan et al. 2006; Morris et al. 2013; Ren et al. 2014). Although having more charged residues, SC16 also has a more distinctive hydrophobic patch. This may allow for efficient alignment and concentration of SC16 at hydrophobic-hydrophilic interfaces, thereby causing faster rodlet assembly and more consistent rodlet morphology (Figs. 2 and 4). The strong dependence of SLH4 assembly on ionic strength suggests that despite having only one charged residue, electrostatic interactions may still be important for its assembly since high salt concentrations reduced assembly.

In conclusion, by comparing the structure and function of two distinct class IB hydrophobins, SC16 and SLH4, we find that these proteins have different assembly morphology and propensity despite having similar tertiary structure and dynamics. SLH4 assembly was most efficient at low ionic strength and a pH of 6.5, whereas SC16 assembly was only modestly influenced by pH and salt with a clear trend of higher pH and higher ionic strength leading to more activity. This difference in assembly propensity may be due to a more localized hydrophobic patch on SC16 providing more robust alignment at interfaces, thus allowing efficient assembly in a wider range of conditions. Circular dichroism spectroscopy indicates that α-helical secondary structure is lost upon rodlet formation, suggesting that in class IB hydrophobins amyloidogenic sequences present in L_1_ undergo a structural change during rodlet formation. Future studies modifying sequences both within and adjacent to L_1_ will further highlight the mechanisms driving self-assembly of these proteins and help to provide rules to select or modify hydrophobins so that they function under specific conditions.

## Acknowledgements

We thank Dr. Mike Lumsden of the Nuclear Magnetic Resonance Research Resource (NMR-3) and Ian Burton from the National Research Council Institute for Marine Biosciences for assistance with NMR data collection. We thank Dr. Jan Rainey for assistance with atomic force microscopy experiments and Dr. David Waisman for providing access to a microplate reader.

## Author Contributions

CK, KLV, and DNL conceived and designed research, conducted experiments, and wrote the manuscript. All authors read and approved the manuscript.

## Funding

This work was funded by the Natural Sciences and Engineering Research Council of Canada (RGPIN-2017-05338). CK was supported by studentships from the Beatrice Hunter Cancer Research Institute and the Dalhousie University Faculty of Medicine. KLV was supported by studentships from the Dalhousie University Faculty of Medicine, the Nova Scotia Health Research Foundation, the NSERC CREATEs BioActives Training Program, researchNS, and the Killam Foundation.

## Data Availability

The authors confirm that the data supporting the findings of this study are available within the article and its supplementary materials.

## Declarations

### Compliance with Ethical Standards

This article does not contain any studies with human participants or animals performed by any of the authors.

### Conflict of Interest

The authors declare no competing interests with the contents of this article.

## Notes

### Competing Interest Statement

The authors have declared no competing interest.

