## Supplementary data for "Characterization of the structure and self-assembly of two distinct class IB hydrophobins"

**Contents**:

Table S1 **Sequences and properties of SC16 and SLH4** 2
Figure S1 **SC16 and SLH4 stabilize oil emulsifications** 3
Figure S2 **NMR resonance assignments of SLH4** 4
Figure S3 **Heteronuclear NOE measurements of SLH4** 5

**Table S1: Sequences and properties of SC16 and SLH4.**

| Protein | Sequence[a] | No. of residues (%) | | pI[b] |
|---|---|---|---|---|
| | | Acidic | Basic | |
| **SC16** | GSTAVPRDVNGGTPPKSCSSGPVYCCNKTED<br>SKHLDKGTTALLGLLNIKIGDLKDLVGLNCS<br>PLSVIGVGGNSCSAQTVCCTNTYQHGLVNVG<br>CTPINIGL | 6 (5.9) | 7 (6.9) | 8.17 |
| **SLH4** | GSGSSQCNAGPVQCCNTLTSASNSQAAGLIQ<br>QLGLSGVGANVPVGINCNPITGIGAGSGSSC<br>NANPACCDNVYTNGLGVQCNPINVNL | 1 (1.2) | 0 (0) | 3.8 |

[a]Residues not native to the hydrophobin are underlined.

[b]Predicted from the protein calculator v.3.4 server; http://protcalc.sourceforge.net. Cysteines were omitted from the sequence during calculation.

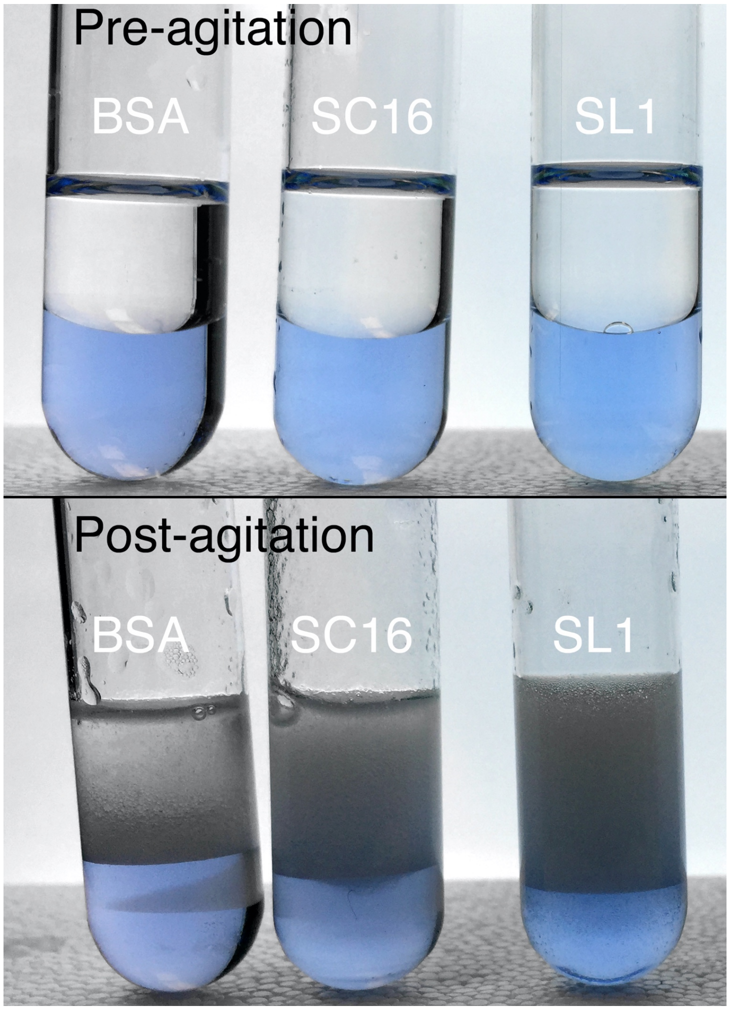


**Figure S1. SC16 and SLH4 stabilize oil emulsifications.** Solutions containing 100 µg/mL BSA, SC16, or SLH4 (dyed blue with 25 µg/mL Remazol brilliant blue for contrast) were vortexed for three minutes with an equal volume of mineral oil. Images are shown of samples pre-agitation and one minute post-agitation.

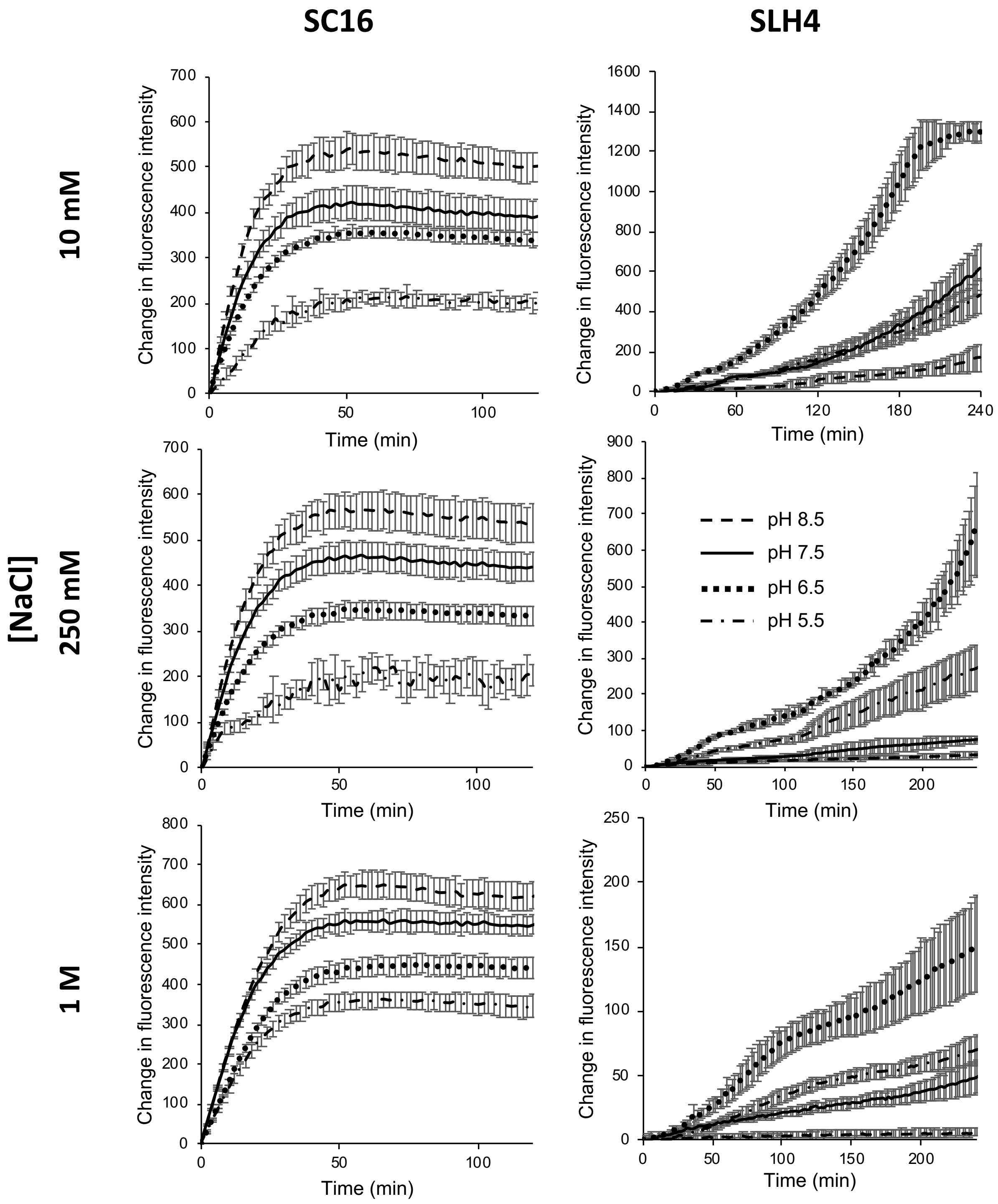


**Figure S2. SC16 and SLH4 self-assembly is influenced by pH and salt**. Thioflavin-T based fluorescence assays were used to quantify rodlet assembly of SC16 and SLH4. Assembly was monitored in solutions with varying pH (5.5-8.5) and at 10 mM, 250 mM or 1 M NaCl. Thioflavin-T fluorescence intensity was monitored at 2 min intervals with 30 s shaking between each interval. Plots show mean + S.D. of 6 replicate experiments.

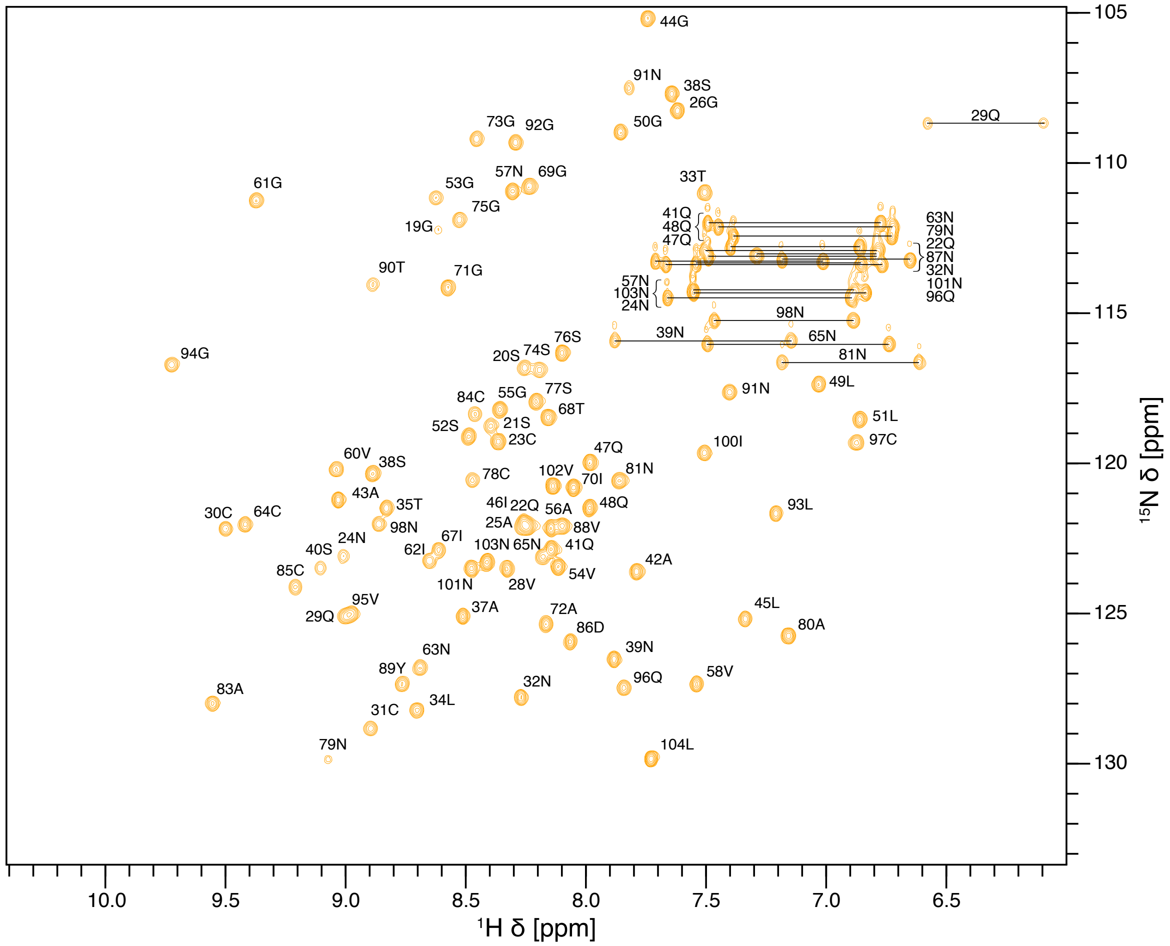


**Figure S3. NMR resonance assignments of SLH4.** The $^{1}$H-$^{15}$N HSQC spectrum of $^{13}$C/$^{15}$N-labelled SLH4 is shown in yellow. Resonance assignments are indicated and peaks corresponding to sidechain amides are connected by a line.

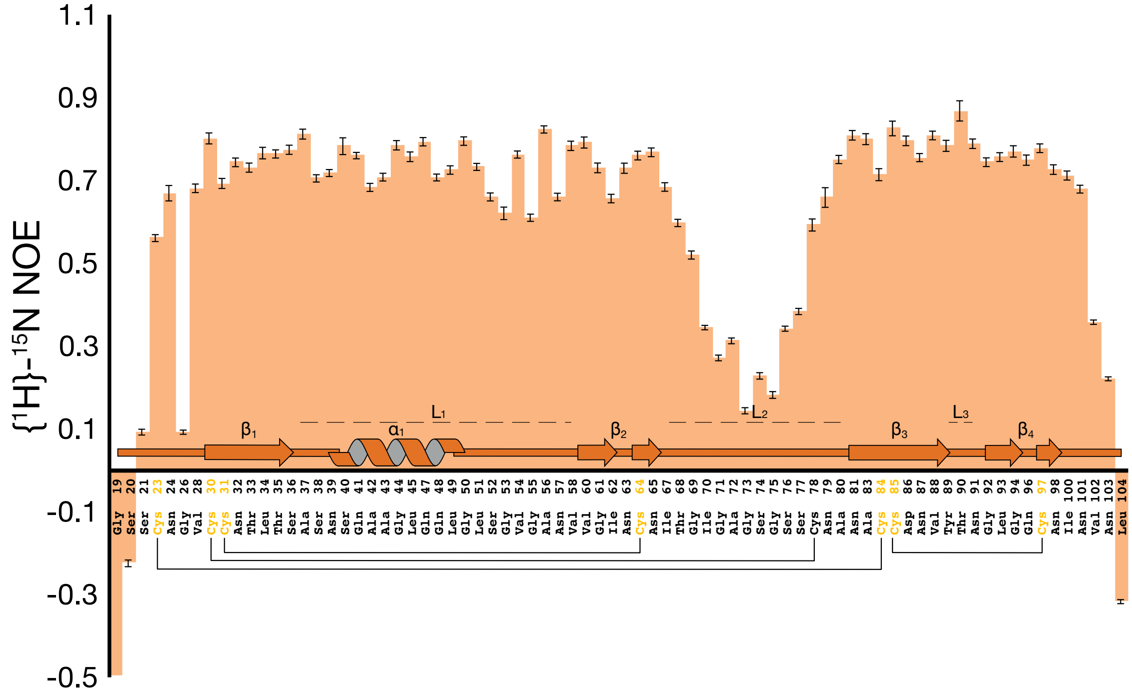


**Figure S4. Heteronuclear NOE measurements of SLH4.** A plot of {$^{1}$H}-$^{15}$N NOE measurements for each residue of SLH4. Secondary structure elements throughout the protein are indicated and conserved disulfide bonds are annotated with lines that connect participating residues.
